# From solitary to colonial with zooid miniaturization: ancestral-state reconstruction based on NGS data of stolidobranch ascidians

**DOI:** 10.1101/2023.12.27.573391

**Authors:** Naohiro Hasegawa, Shin Matsubara, Akira Shiraishi, Honoo Satake, Noa Shenkar, Hiroshi Kajihara

## Abstract

The size of organisms has consistently intrigued researchers across various disciplines in biology. However, the evolutionary process of zooid miniaturization in colonial animals remained an enigmatic topic. The family Styelidae, within the ascidian order Stolidobranchia, showcases a diverse spectrum of coloniality, positioning it as an ideal candidate for delving into the intricacies of colonial evolution. In this research, we inferred a phylogenomic relationship mainly within Styelidae using transcriptomes of a total of 42 ascidians; from 17 species sampled in Israel and Japan and transcriptome data from 25 species sourced from a previous study and a database. Through ancestral-state reconstruction, our analysis indicated a clear directional change: following the acquisition of coloniality, zooids tended to become progressively smaller. This miniaturization is likely an adaptive response, enabling organisms to swiftly colonize limited marine substrate. We formulated a mathematical model suggesting that zooid miniaturization, due to living space constraints, would result in a faster asexual cycle and accelerated expansion in a colony. Our data also suggested that coloniality evolved independently three times within Styelidae. Moreover, once colonial traits are established, they appear to be consistently preserved, underscoring their biological importance in the colonial lineage.

## Introduction

The body size of organisms has been a matter of research topic among biologists, with its related factors so far considered including latitudinal variations, physiological implications, and life history strategies (Grischenko et al. 2002). Yet, much of our understanding of the evolutionary shifts in animal body size is derived primarily from solitary metazoans rather than colonial ones (Liow and Taylor 2019).

Colonies of sessile aquatic animals consist of modular individuals, termed polyps and zooids (Beklemishev 1969). The modules within a single colony are clones, proliferating through asexual reproduction (Ryland and Warner 1986). Coloniality has independently evolved across various animal higher taxa (Hiebert et al. 2021), including Bryozoa (Schwaha et al. 2020), Cnidaria (Barbeitos et al. 2010; Kayal et al. 2018), Hemichordata (Cannon et al. 2013), Kamptozoa (Fuchs et al. 2010), and Tunicata (Alié et al. 2021). Within each taxon, the modules of colonial organisms are typically smaller than solitary individuals (Ryland and Warner 1986; Mukai 2001). Although zooid miniaturization has been noticed in various taxa for approximately 40 years (e.g., Kott 1985; Ryland and Warner 1986; Davidson et al. 2004; Sogot et al. 2014; Kocot 2016; Braun and Stach 2018; Nanglu et al. 2023), its evolutionary process still remains a topic of inquiry.

Ascidians, commonly known as sea squirts, exhibit a wide range of life history strategies (Satoh 1994; Burighel and Cloney 1997; Davidson et al. 2004). Within Ascidiacea, the order Stolidobranchia has been recognized as an ideal group for exploring the evolutionary plasticity of coloniality (Zeng et al. 2006; Pérez-Portela et al. 2009; Brown and Swalla 2012; Alié et al 2021). This order encompasses three families, Molgulidae, Pyuridae, and Styelidae (cf. Kott 1985; Monniot et al. 1991). While both molgulid and pyurid ascidians are solitary, styelids comprise solitary and colonial species (cf. Kott 1985; Monniot et al. 1991). Kott (1985) classified Styelidae into three subfamilies: Botryllinae, which includes colonial forms with systems—zooids’ arrangements around common cloacal cavities in a colony; Polyzoinae, which comprises colonial forms without systems; and Styelinae, which consists solely of solitary forms.

Among the colonial groups, a diverse range of coloniality has been observed (Fig. 1). In terms of the degree of colonial complexity, the polyzoine genus *Symplegma* Herdman, 1886 positions between other polyzoines and the highly integrated botryllines (Shirae et al. 1999; Gutierrez and Brown 2017). There is no morphological difference between the colonial genus *Syncarpa* Redikorzev, 1913 and the solitary genus *Dendrodoa* MacLeay, 1824 except forming colonies (Hasegawa and Kajihara 2019). The diversity of coloniality in Styelidae is actualized by various budding modes, which can be categorized based on the tissues from which buds (= daughter zooids) originate in mother zooids. Specifically, three budding modes are recognized: vasal budding (Alié et al. 2018), peribranchial budding (Metschnikoff 1869; Pizon 1893; Ritter 1896; SelysLLongchamps 1917; Berrill 1940, 1948; Watanabe and Tokioka 1972; Mukai and Watanabe 1976; Kawamura and Watanabe 1981; Akhmadieva et al. 2007), and vascular budding (Giard 1872; Oka and Watanabe 1957; Sabbadin et al. 1975; Nakauchi 1982; Brunetti and Mastrototaro 2004; Brown et al. 2009; Kawamura and Sunanaga 2010; Gutierrez and Brown 2017).

**Figure 1.**
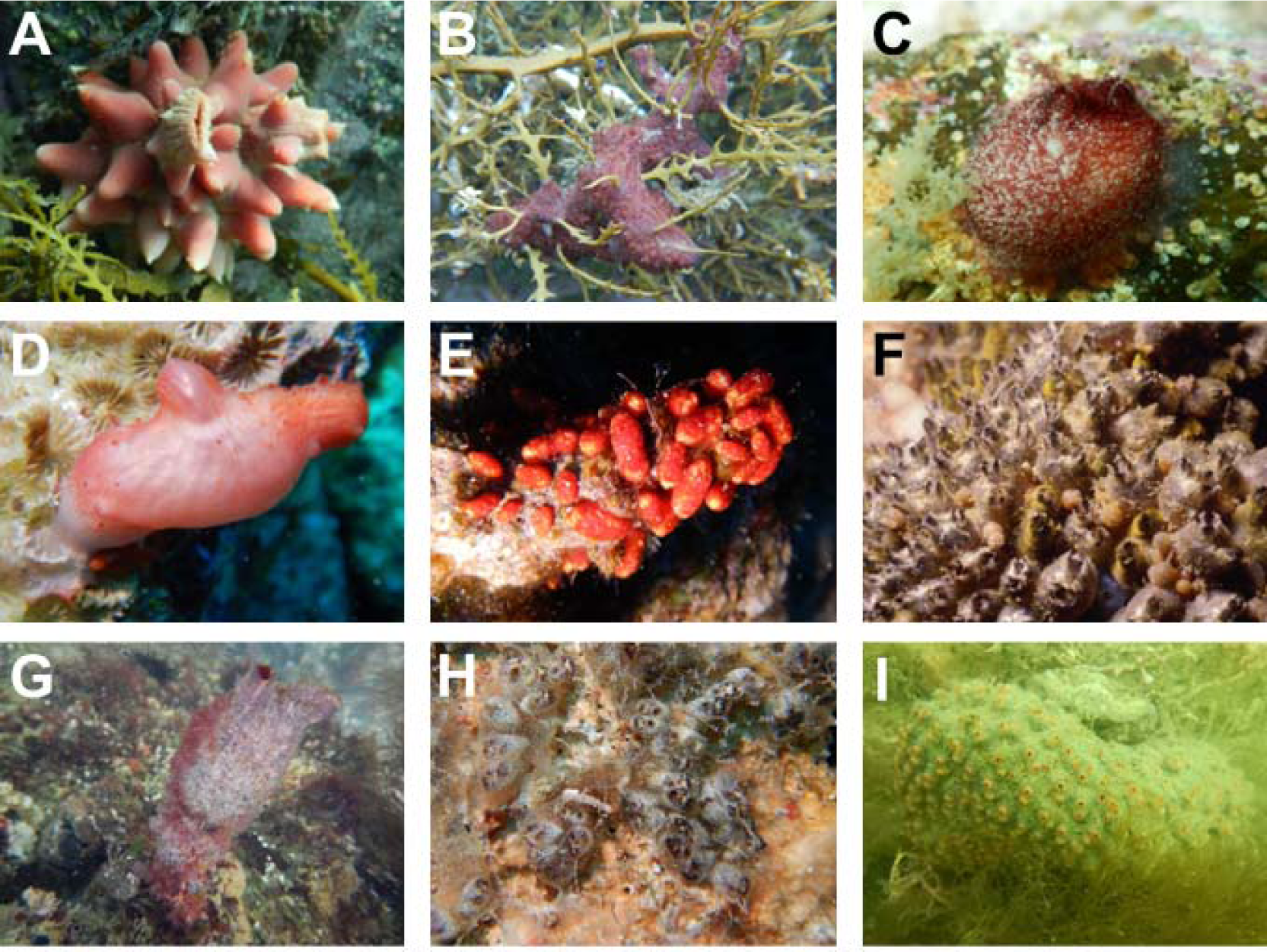
Solitary and colonial ascidians in Stolidobranchia. A, *Halocynthia roretzi*; B, *Botrylloides violaceus*; C, *Cnemidocarpa clara*; D, *Cnemidocarpa margaritifera*; E, *Eusynstyela latericius*; F, *Polyandrocarpa zorritensis* (photographed by Dr. Nishikawa); G, *Styela clava*; H; *Symplegma systematica*; I, *Syncarpa composita*.

Phylogenetic analyses of Stolidobranchia based on partial sequences of 18S rRNA and mitochondrial cytochrome *c* oxidase subunit I genes have variably suggested that coloniality evolved once (Zeng et al. 2006; Tsagkogeorga et al. 2009; Hasegawa and Kajihara 2019) or seven times (Pérez-Portela et al. 2009), but these studies often yielded trees with less-supported nodes. Alié et al. (2018) suggested that peribranchial budding and vasal budding evolved convergently, and that vascular budding was secondarily derived in an ancestor exhibiting peribranchial budding, based on a phylogenomic tree with high support values. However, their analysis (Alié et al. 2018) lacked some key taxa, such as *Symplegma* and *Syncarpa*, which are pivotal for discussions on the evolution of coloniality. While the phylogenetic relationships in Stolidobranchia have been inferred by many previous authors to elucidate how coloniality evolved, no attempt has been made to reconstruct the ancestral states of coloniality using statistical methods.

The aim of this study is to clarify the evolutionary relationship between zooid size and coloniality. A robust phylogenomic tree was constructed, encompassing key taxa in the Stolidobranchia essential for understanding the evolution of coloniality. Through Bayesian inference for ancestral-state reconstruction, we provide a clear depiction of trait evolution related to coloniality on the tree. Additionally, we propose a mathematical model suggesting that zooid miniaturization has been driven by spatial competition on substrates.

## Methods

### Sampling, fixation, and identification

A total of 17 ascidian species, including both colonial and solitary forms, were sampled in Israel and Japan from May 2021 to January 2023 (Table 1). We conducted taxon sampling without bias in body size. The animals were anesthetized with menthol. For solitary individuals, oral or atrial siphons were excised and preserved in RNAlater; the remaining bodies were preserved in 10% formalin. For colonial forms, a part of the colonies was cut off and preserved in RNAlater; another part was preserved in 10% formalin. The RNAlater-preserved specimens were used for total RNA extraction; the formalin-fixed specimens were used for morphological identification. For specimens collected in Israel, 1% borax was added to the formalin fixative solution.

**Table 1.** Sample information including order, family, species name, sampling date, and sampling site.

| Order | Family | Species | Sampling date | Sampling site |
| --- | --- | --- | --- | --- |
| Aplousobranchia | Clavelinidae | *Clavelina lepadiformis* (Müller, 1776) | 2021/5/12 | Kushimoto, Japan |
|  | Polyclinidae | *Aplidium* sp. | 2022/3/9 | Misaki, Japan |
|  | Pseudodistomidae | *Pseudodistoma antinboja* Tokioka, 1949 | 2021/5/11 | Koza, Japan |
| Stolidobranchia | Pyuridae | *Herdmania momus* (Savignyi, 1816) | 2021/5/11 | Koza, Japan |
|  |  | *Microcosmus* sp. | 2021/5/11 | Koza, Japan |
|  |  | *Pyura mirabilis* (Heller, 1877) | 2021/11/1 | Sugashima Island, Japan |
|  | Styelidae | *Botrylloides* sp. | 2022/3/15 | Misaki, Japan |
|  |  | *Botrylloides violaceus* Oka, 1927 | 2021/11/1 | Sugashima Island, Japan |
|  |  | *Cnemidocarpa clara* (Hartmeyer, 1906) | 2021/11/1 | Sugashima Island, Japan |
|  |  | *Cnemidocarpa margaritifera* Michaelsen, 1918 | 2023/1/18 | Eilat, Israel |
|  |  | *Eusynstyela latericius* (Sluiter, 1904) | 2023/1/16 | Eilat, Israel |
|  |  | *Eusynstyela misakiensi* (Watanabe and Tokioka, 1972) | 2021/5/11 | Koza, Japan |
|  |  | *Polycarpa cryptocarpa kroboja* (Sluiter, 1885) | 2021/11/1 | Sugashima Island, Japan |
|  |  | *Dendrodoa aggregate* (Rathke, 1806) | 2022/6/10 | Akkeshi, Japan |
|  |  | *Styela* sp. | 2022/3/15 | Misaki, Japan |
|  |  | *Symplegma systematica* (Sluiter, 1904) | 2022/4/23 | Irabujima Island, Japan |
|  |  | *Syncarpa composita* (Tokioka, 1951) | 2022/6/30 | Akkeshi, Japan |

### RNA extraction, purification, and sequencing

The tissues preserved in RNAlater were homogenized in tubes containing 500 mL of Sepazol-RNA I Super G (Nacalai tesque, Kyoto). The homogenate was left at room temperature for 5 minutes and then mixed with 100 μL of chloroform. Then, the samples were centrifuged at 12,000 × g for 15 minutes. 250 μL of upper aqueous phase was mixed with 250 μL of isopropanol and 0.5 μL of glycogen solution, followed by incubation at room temperature for 10 minutes. Subsequently, they were centrifuged at 12,000 × g for 10 minutes. The solution was removed from tubes, and 500 μL of 80% ethanol was added. After centrifugation at 12,000 × g for 5 minutes, the ethanol was carefully removed; then, pellets were air-dried. The pellets were dissolved in 200 μL of TE to obtain crude RNA extracts. The crude RNA solution (dissolved in 200 μL of TE) was added to 200 μL of a mixture of phenol:chloroform:isoamyl alcohol (25:24:1). The samples were centrifuged at 13,000 rpm for 10 minutes at room temperature. Upper aqueous phase was transferred to a new tube; then, 200 μL of chloroform:isoamyl alcohol (24:1) was added. Once more, after centrifugation at 13,000 rpm for 10 minutes at room temperature, 180 μL of upper aqueous phase was collected. To this, 20 μL of 3M sodium acetate solution, 500 μL of 100% ethanol, and 0.5 μL of glycogen solution were added. The samples were then centrifuged at 15,000 rpm for 30 minutes at 4°C, and the supernatant was removed. Pellets were mixed with 700 μL of 80% ethanol, followed by centrifugation at 15,000 rpm for 5 minutes at 4°C. The supernatant was removed, and pellets were air-dried. The pellets were dissolved in 10 μL of TE to obtain total RNA solutions. Concentration of total RNA in the solution was measured using a NanoDrop (Thermo Fisher Scientific). The volume of the solution containing 5 μg of RNA was measured; then, pure water was added to adjust the total volume to 26.5 μL; for samples that did not meet the 5 μg threshold, the entire volume of the total RNA solution was added. For removing DNA from the samples, 3 μL of 10 × TURBO DNase buffer (Thermo Fisher Scientific) and 0.5 μL of TURBO DNase (Thermo Fisher Scientific) were added. The samples were incubated at 37°C for 30 minutes using a 2720 thermal cycler (Thermo Fisher Scientific). After incubation, 3 μL of DNase Inactivation Reagent (Thermo Fisher Scientific) was added, and the samples were incubated at room temperature for 5 minutes with tapping every 2 min. Subsequently, the samples were centrifuged at 10,000 × g for 2 minutes at room temperature, and 25 μL of supernatant was collected. To the supernatant, 75 μL of water, 10 μL of 3M sodium acetate solution, 0.5 μL of glycogen solution, and 275 μL of 100% ethanol were added. The samples were centrifuged at 15,000 rpm for 30 minutes at 4°C; then, the supernatant was removed. Pellets were mixed with 400 μL of 80% ethanol, followed by centrifugation at 15,000 rpm for 5 minutes at 4°C. Once more, supernatant was removed; the pellets were air-dried. The pellets were dissolved in 11 μL of pure water to obtain total RNA solutions free of genomic DNA. Concentration of RNA in the solutions was measured using a NanoDrop (Thermo Fisher Scientific). Absence of RNA degradation was confirmed by electrophoresis. cDNA library preparation and paired-end sequencing were performed by Novogene; the Illumina NovaSeq 6000 was used for next-generation sequencing.

### Phylogenomic analysis

For obtaining *de novo* transcriptomes, we mainly followed Alié et al. (2018). The following three types of Illumina reads were filtered by Novogene: reads containing adapter contamination, with more than 10% of bases that could not be determined throughout the entire length, and where more than 50% of bases had Q20 of Phred value (Ewing et al. 1998) in their entire length. The reads data was registered as the BioProject PRJNA880804 in Sequnce Read Archive (SRA). The filtered reads were *de novo* assembled with Trinity ver. 2.8.5 (Grabherr et al. 2011) using default parameters. Open reading frames (ORFs) with less than 90 amino acids were discarded by TransDecoder ver. 5.5.0 (https://github.com/TransDecoder/TransDecoder; last accessed May 2023). ORFs having similar nucleotide sequences were clustered using CD-HIT-EST ver. 4.8.1 (Li and Godzik 2006; Fu et al. 2012) with default parameters. The reads were mapped on the remaining contigs by using Kallisto ver. 0.46.2 (Bray et al. 2016). The contigs that had low expression levels were picked up by drawing density plots with R; the contigs were retained based on the threshold corresponding to the lower peak of the plot. To assess the accuracy of the sequencing, BUSCO ver. 5.3.2 (Simão et al. 2015; Waterhouse et al. 2017; Seppey et al. 2019; Manni et al. 2021) with metazoan datasets was used to examine the presence of core gene sets in the assembled sequences. Cross-contamination occurring in the process from the extraction to the RNA-seq was removed using CroCo (Simion et al. 2018). In addition to the *de novo* transcriptomes, 16 transcriptomes were downloaded from a GitHub repository, styelidae (Alié et al. 2018; https://github.com/AlexAlie/styelidae) for a phylogenomic analysis (Table 2); core proteomes of nine ascidian species were also downloaded from Ascidian Network for In Situ Expression and Embryological Data (ANISEED) (Table 2). I retained only the longest sequence for each gene in their respective proteomes from each of the nine species; CD-HIT ver. 4.8.1 (Fu et al. 2012) was used for dereplicating with -c *1.0* option, for excluding sequences consisting of less than 30 amino acids.

**Table 2.**
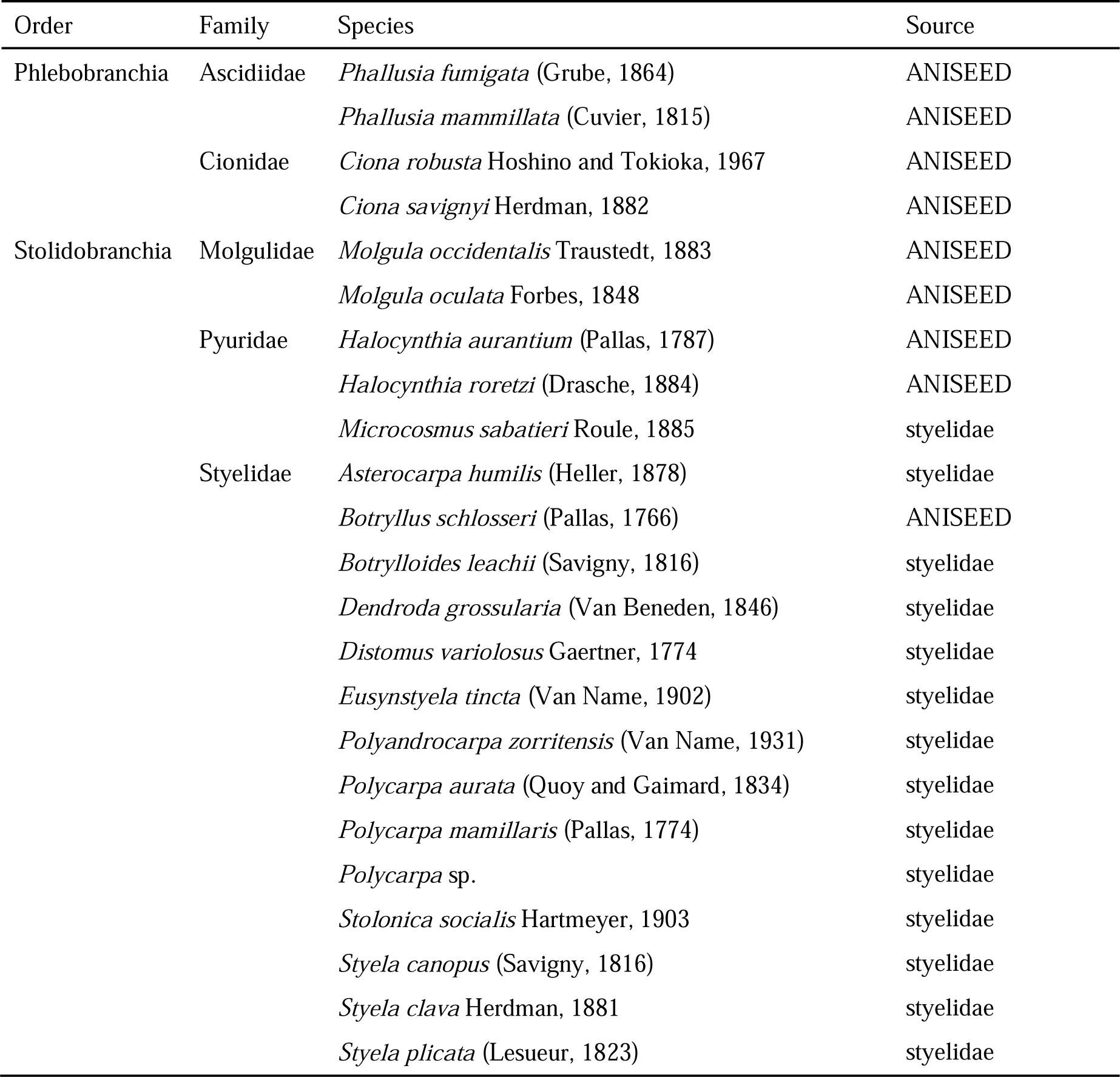
Sequence data obtained from the GitHub repository styelidae and ANISEED.

Multiple alignments were constructed for the phylogenomic analysis. Orthologous clusters were built based on the proteomes from 42 species using OrthoFinder ver. 2.5.4 (Emms and Kelly 2015) with -M *msa* and -S *blast* options. The sequences were downsampled within each cluster that had multiple sequences derived from the same sample; the longest sequence was designated as the representative sequence for each species. From these downsampled clusters, we excluded the clusters that did not meet the following criteria: within each cluster, *i*) containing 30 or more sequences of the total 33 species listed in Table 1 and from the GitHub repository styelidae (Table 2); and *ii*) containing seven or more sequences of the nine species from ANISEED (Table 2). For these two steps, we utilized a Python script in the GitHub repository Directionality_zooid_miniaturization (https://github.com/231007NHasegawa/Directionality_zooid_miniaturization). Each remained cluster was aligned by MAFFT ver. 7.520 (Katoh and Standley 2013) with *E-INS-i* strategy. The alignments were individually trimmed using TrimAl ver. 1.4. rev. 15 (Capella-Gutiérrez et al. 2009) with *strictplus* method. Then, the trimmed alignments were concatenated with each other by my original Python script (https://github.com/231007NHasegawa/Directionality_zooid_miniaturization).

Maximum Likelihood (ML) analysis and Bayesian inference (BI) were performed based on the concatenated alignment for inferring a phylogenetic relationship among Stolidobranchia. Prior to conducting the ML analysis, the substitution models corresponding to each partition in the multiple alignment were selected with ModelFinder program (Kalyaanamoorthy et al. 2017) referring partition models (Chernomor et al. 2016) implemented in IQ-TREE ver. 2.2.2.6 (Minh et al. 2020); then, the ML analysis was conducted by IQ-TREE ver. 2.2.2.6 (Minh et al. 2020) with ultrafast (UF) bootstrap method (Hoang et al. 2018). Branch support values were evaluated with 1,000 UF bootstrap replicates. BI analysis was performed using the MPI version of ExaBayes ver. 1.5.1 (Aberer et al. 2014). Two independent MCMC runs were parallelly run, started from a random tree; cold chain and heated chain with 0.5 of “heatFactor” option were executed per independent run. The runs were stopped that each chain was run for 500,000 generations; at that time, the average standard deviation of split frequencies reached 0.0%. Trees were sampled every 100 generations; the initial 25% of sampled trees were discarded as burn-in. Convergence was checked by postProcParam program implemented in ExaBayes.

### Ancestral-state reconstruction

To determine values for matrix entries in transition probability matrices for each character—*i*) vasal budding, *ii*) peribranchial budding, *iii*) vascular budding, *iv*) common cloacal cavity, and *v*) coloniality (colonial or solitary)—we initially computed a maximum likelihood estimate by using “MultiState” (Pagel et al. 2004) as implemented in BayesTraits ver. 4.0.1 (http://www.evolution.reading.ac.uk/BayesTraitsV4.0.1/BayesTraitsV4.0.1.html; last accessed September 30th, 2023) for a transition rate of each character state. The rates calculated as zero were simply adjusted to 0.01 in the matrices, indicating that such transitions were rare evolutionary events. Subsequently, we conducted Bayesian inferences (BI) using “MultiState” (Pagel et al. 2004) based on the ML tree with 10,000,000 iterations, a burn-in of 2,500,000, a sampling frequency of 1,000, and the “ScaleTrees” option set at 0.1 because a mean of branch lengths of the ML tree was ca. 0.1. I employed a uniform distribution as a non-informative prior, adjusting an interval such that the transition probability matched the mean of the uniform distribution. Expected a posteriori (EAP) was computed to represent the posterior probabilities (PPs) of the character states at each node.

For the ancestral-state reconstruction, we obtained literature-based data of the longest body length for each species based on taxonomic descriptions (Table 3). For *Botrylloides* sp., *Microcosmus* sp., and *Styela* sp., the body lengths were measured based on the formalin-fixed specimens. Information about the body length of *Polycarpa* sp. is lacking (Alié et al. 2018). Based on the phylogenomic tree, MCMC analysis was conducted selecting Brownian motion model by “Continuous: Random Walk” (Pagel 1999) with 10^8^ iterations, 25% of burn-in, and the “ScaleTrees” option set 0.1. The body length of each node was represented by a mean of values sampled from every 1,000 generations.

**Table 3.**
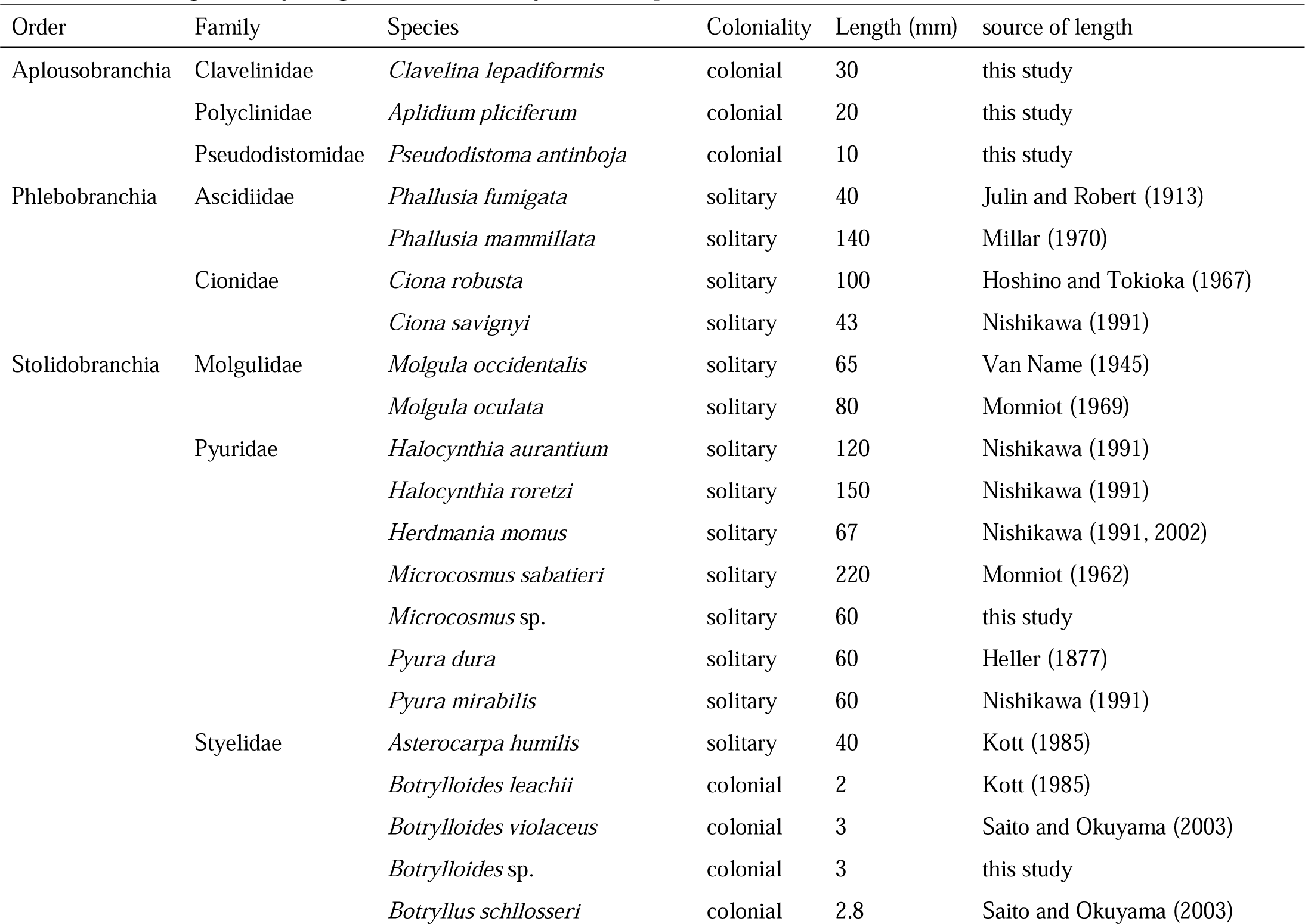
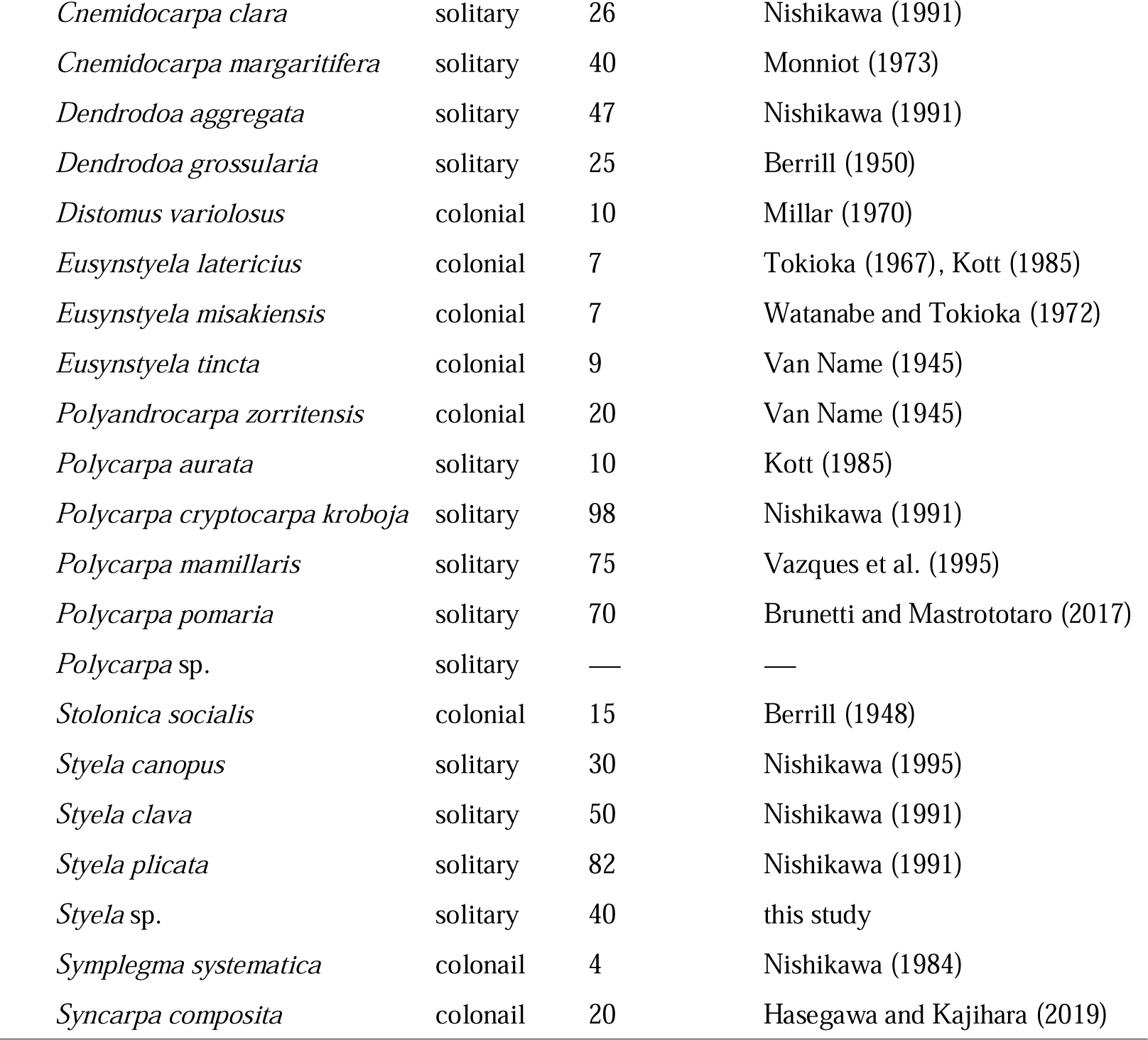
The longest body length and coloniality of each species1.

| Order | Family | Species | Coloniality | Length (mm) | source of length |
| --- | --- | --- | --- | --- | --- |
| Aplousobranchia | Clavelinidae | *Clavelina lepadiformis* | colonial | 30 | this study |
|  | Polyclinidae | *Aplidium pliciferum* | colonial | 20 | this study |
|  | Pseudodistomidae | *Pseudodistoma antinboja* | colonial | 10 | this study |
| Phlebobranchia | Asciidiidae | *Phallusia fumigata* | solitary | 40 | Julin and Robert (1913) |
|  |  | *Phallusia mammillata* | solitary | 140 | Millar (1970) |
|  | Cionidae | *Ciona robusta* | solitary | 100 | Hoshino and Tokioka (1967) |
|  |  | *Ciona savignyi* | solitary | 43 | Nishikawa (1991) |
| Stolidobranchia | Molgulidae | *Molgula occidentalis* | solitary | 65 | Van Name (1945) |
|  |  | *Molgula oculata* | solitary | 80 | Monniot (1969) |
|  | Pyuridae | *Halocynthia aurantium* | solitary | 120 | Nishikawa (1991) |
|  |  | *Halocynthia roretzi* | solitary | 150 | Nishikawa (1991) |
|  |  | *Herdmania momus* | solitary | 67 | Nishikawa (1991, 2002) |
|  |  | *Microcosmus sabatieri* | solitary | 220 | Monniot (1962) |
|  |  | *Microcosmus* sp. | solitary | 60 | this study |
|  |  | *Pyura dura* | solitary | 60 | Heller (1877) |
|  |  | *Pyura mirabilis* | solitary | 60 | Nishikawa (1991) |
|  | Styelidae | *Asterocarpa humilis* | solitary | 40 | Kott (1985) |
|  |  | *Botrylloides leachii* | colonial | 2 | Kott (1985) |
|  |  | *Botrylloides violaceus* | colonial | 3 | Saito and Okuyama (2003) |
|  |  | *Botrylloides* sp. | colonial | 3 | this study |
|  |  | *Botryllus schlosseri* | colonial | 2.8 | Saito and Okuyama (2003) |

|  |  |  |  |
| --- | --- | --- | --- |
| *Cnemidocarpa clara* | solitary | 26 | Nishikawa (1991) |
| *Cnemidocarpa margaritifera* | solitary | 40 | Monniot (1973) |
| *Dendrodoa aggregata* | solitary | 47 | Nishikawa (1991) |
| *Dendrodoa grossularia* | solitary | 25 | Berrill (1950) |
| *Distomus variolosus* | colonial | 10 | Millar (1970) |
| *Eusynstyela latericius* | colonial | 7 | Tokioka (1967), Kott (1985) |
| *Eusynstyela misakiensis* | colonial | 7 | Watanabe and Tokioka (1972) |
| *Eusynstyela tincta* | colonial | 9 | Van Name (1945) |
| *Polyandrocarpa zorritensis* | colonial | 20 | Van Name (1945) |
| *Polycarpa aurata* | solitary | 10 | Kott (1985) |
| *Polycarpa cryptocarpa kroboja* | solitary | 98 | Nishikawa (1991) |
| *Polycarpa mamillaris* | solitary | 75 | Vazques et al. (1995) |
| *Polycarpa pomaria* | solitary | 70 | Brunetti and Mastrototaro (2017) |
| *Polycarpa* sp. | solitary | — | — |
| *Stolonica socialis* | colonial | 15 | Berrill (1948) |
| *Styela canopus* | solitary | 30 | Nishikawa (1995) |
| *Styela clava* | solitary | 50 | Nishikawa (1991) |
| *Styela plicata* | solitary | 82 | Nishikawa (1991) |
| *Styela* sp. | solitary | 40 | this study |
| *Symplegma systematica* | colonail | 4 | Nishikawa (1984) |
| *Syncarpa composita* | colonail | 20 | Hasegawa and Kajihara (2019) |

### Character correlation analysis

To test the correlation between coloniality and body length reconstructed on the tree nodes (whether or not there is any evolutionary tendency between these characters), we utilized the “Test trait correlations: continuous” and “Independent Contrast: Correlation” (Felsenstein 1973; Freckleton 2012) in BayesTraits version 4.0.1. Both MCMC approaches were adopted, employing a stones command, setting 100 stones with 1,000 iterations each to determine a marginal likelihood. Subsequent analyses were implemented using the “TestCorrel” command, which sets the covariance to zero, employing identical command parameters as the first analysis. Log Bayes Factor (Log BF) was calculated by taking twice the difference of log marginal likelihoods between the two analyses.

### Detection of evolutionary direction

The resulting tree included a subclade solely consisting of members that undergo peribranchial budding (= Clade A, see below). To evaluate if the body size evolved directionally in the entire Stolidobranchia and this subclade, respectively, MCMC analyses were separately conducted based on the whole of the ML tree and this subclade. First, an MCMC analysis was executed with the command selecting “Continuous: Random Walk (Model A)” (Pagel 1999) and a stones command estimating a marginal likelihood using 100 stones and 1000 iterations per stone. Second, another MCMC analysis was carried out with the “Continuous: Directional (Model B)” method (Pagel 1999), maintaining the same parameters for the stones command. Then, the Log BF was computed by subtracting the log marginal likelihood of the Model A and Model B. In addition, we calculated correlation coefficient between body length and branch length for solitary and colonial forms.

## Results

### Phylogenomics

The number of Illumina reads after filtration obtained from 17 species ranged from 42 million to 137 million. Through the assembly process, 26,116–108,633 contigs were kept for each species (Table 4). These contigs, along with transcriptomes of 16 species from Alié et al. (2018) and proteomes of nine species from ANISEED, were clustered in 360,094 orthologous groups. The number of orthologous groups that met the filtration criteria were 1,883 out of 360,094. After alignment, trimming, and concatenation, the dataset comprised 1,039,648 aa including gaps of 42 ascidian species with 0.1% of missing data. The dataset was partitioned into 130 blocks; the best-fit model was selected in each partition.

**Table 4.** Number of reads and contigs of each species among each step of quality control.

| Species | Number of reads | Trinity | Transdecoder | CD-HIT | min TPM | CroCo | ratio | Busco Index (%) |
| --- | --- | --- | --- | --- | --- | --- | --- | --- |
| *Clavelina lepadiformis* | 56682322 | 126660 | 122351 | 36843 | 36843 | 36843 | 29.1 | 96.3 |
| *Aplidium pliciferum* | 55774938 | 115622 | 95490 | 39219 | 39219 | 39219 | 33.9 | 94.1 |
| *Pseudodistoma antinboja* | 61465810 | 253926 | 130804 | 50694 | 50694 | 50694 | 20.0 | 94.9 |
| *Herdmania momus* | 42434406 | 88008 | 89631 | 26116 | 26116 | 26116 | 29.7 | 94 |
| *Microcosmus* sp. | 54163154 | 87164 | 106927 | 30896 | 30896 | 30896 | 35.4 | 95.6 |
| *Pyura mirabilis* | 43199584 | 83141 | 89898 | 29224 | 29224 | 29224 | 35.1 | 92.5 |
| *Botrylloides* sp. | 137362636 | 201787 | 137979 | 65432 | 65432 | 65432 | 32.4 | 96 |
| *Botrylloides violaceus* | 122384370 | 147738 | 112922 | 49286 | 49286 | 49286 | 33.4 | 95.8 |
| *Cnemidocarpa clara* | 43854848 | 114765 | 96283 | 34223 | 34223 | 34223 | 29.8 | 93.4 |
| *Cnemidocarpa margaritifera* | 44546458 | 72124 | 89219 | 27893 | 27893 | 27893 | 38.7 | 94.4 |
| *Eusynstyela latericius* | 124303040 | 344331 | 146190 | 74497 | 74497 | 74497 | 21.6 | 92.8 |
| *Eusynstyela misakiensi* | 127341340 | 327188 | 265280 | 108633 | 108633 | 108633 | 33.2 | 99.2 |
| *Polycarpa cryptocarpa kroboja* | 51953554 | 130749 | 122330 | 46481 | 46481 | 46481 | 35.5 | 95.3 |
| *Dendrodoa aggregata* | 46787562 | 172013 | 157965 | 71393 | 71393 | 71393 | 41.5 | 97.4 |
| *Styela* sp. | 56036346 | 140726 | 120963 | 43202 | 43202 | 43202 | 30.7 | 96.5 |
| *Symplegma systematica* | 126899150 | 314471 | 126874 | 64540 | 64540 | 64540 | 20.5 | 93.4 |
| *Syncarpa composita* | 134010390 | 501552 | 229526 | 99555 | 99555 | 99555 | 19.8 | 94.4 |

Almost all the nodes in the tree received highly supported values (100% UF bootstrap; 1.00 PP); as an exception, the clade of *Eusynstyela tincta* + *E. latericius* + *E. misakiensis* has 85% UF bootstrap value (Fig. 2). The order Stolidobranchia was sister to the clade Aplousobranchia + Phlebobranchia. Phlebobranchia was non-monophyletic; in it, *Phallusia* was sister to Aplousobranchia. In Aplousobranchia, *Aplidium* formed a clade with *Pseudodistoma*. In Stolidobranchia, Molgulidae was inferred to have branched off first from the rest of this group; Styelidae was recovered as monophyletic, which turned out to be sister to a part of paraphyletic Pyuridae. In Styelidae, members that perform peribranchial budding—botryllines and part of polyzoines—formed a clade (hereafter Clade A); in Clade A, *Symplegma* was sister to Botryllinae (*Botrylloides* and *Botryllus*). *Syncarpa* grouped together with paraphyletic *Dendrodoa*.

**Figure 2.**
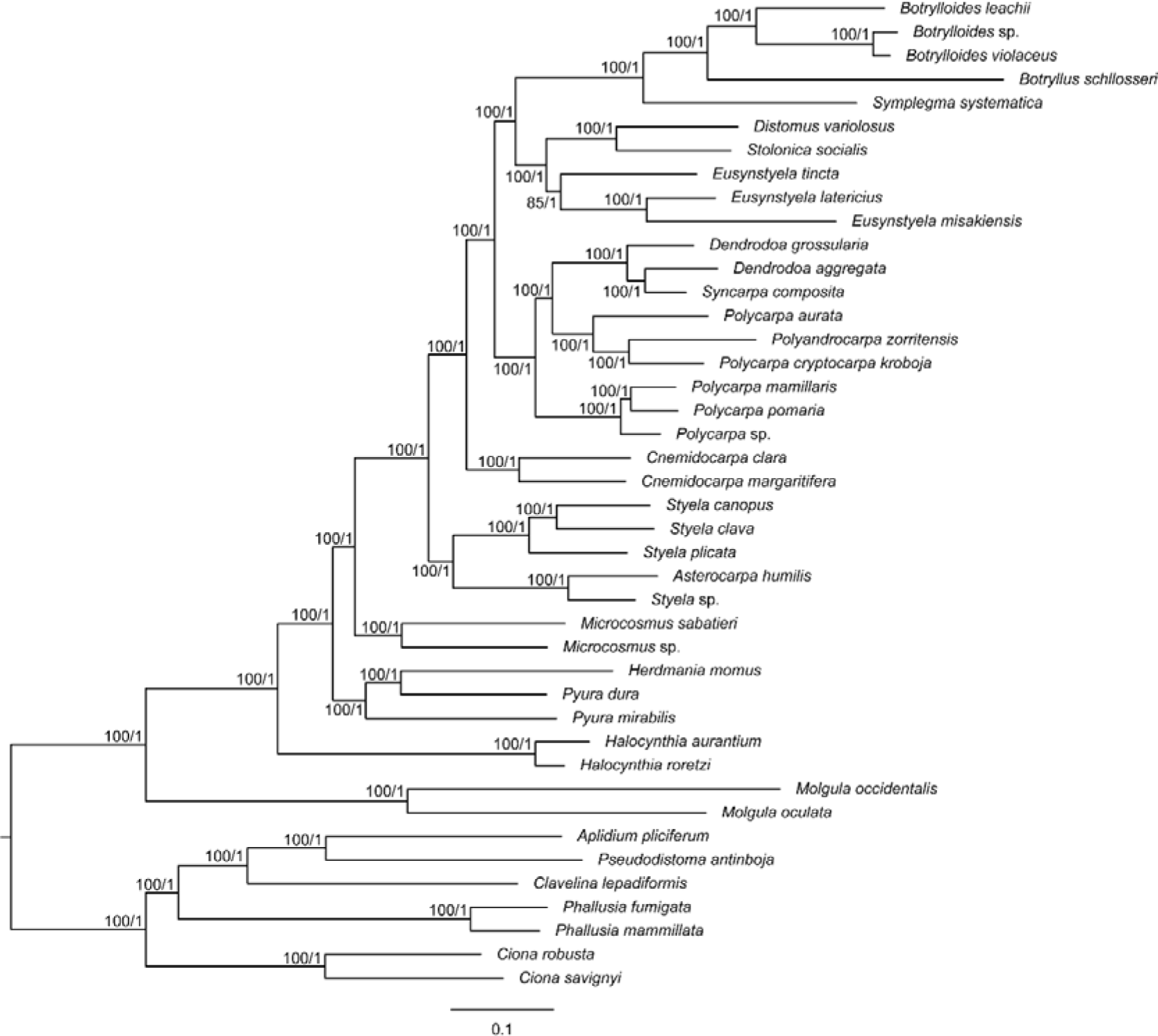
Maximum-likelihood tree based on 1,883 genes; the number on each branch indicates UF bootstrap value and PP.

### Ancestral-state reconstruction

The EAPs for the coloniality-related character states—either *i*) budding from vasal stolon, *ii*) budding from peribranchial epithelium, *iii*) budding from blood vessel, *iv*) with a common cloacal cavity, or *v*) non colonial (i.e., solitary)—exceeded 99% in every node (Figs 3–7).

**Figure 3.**
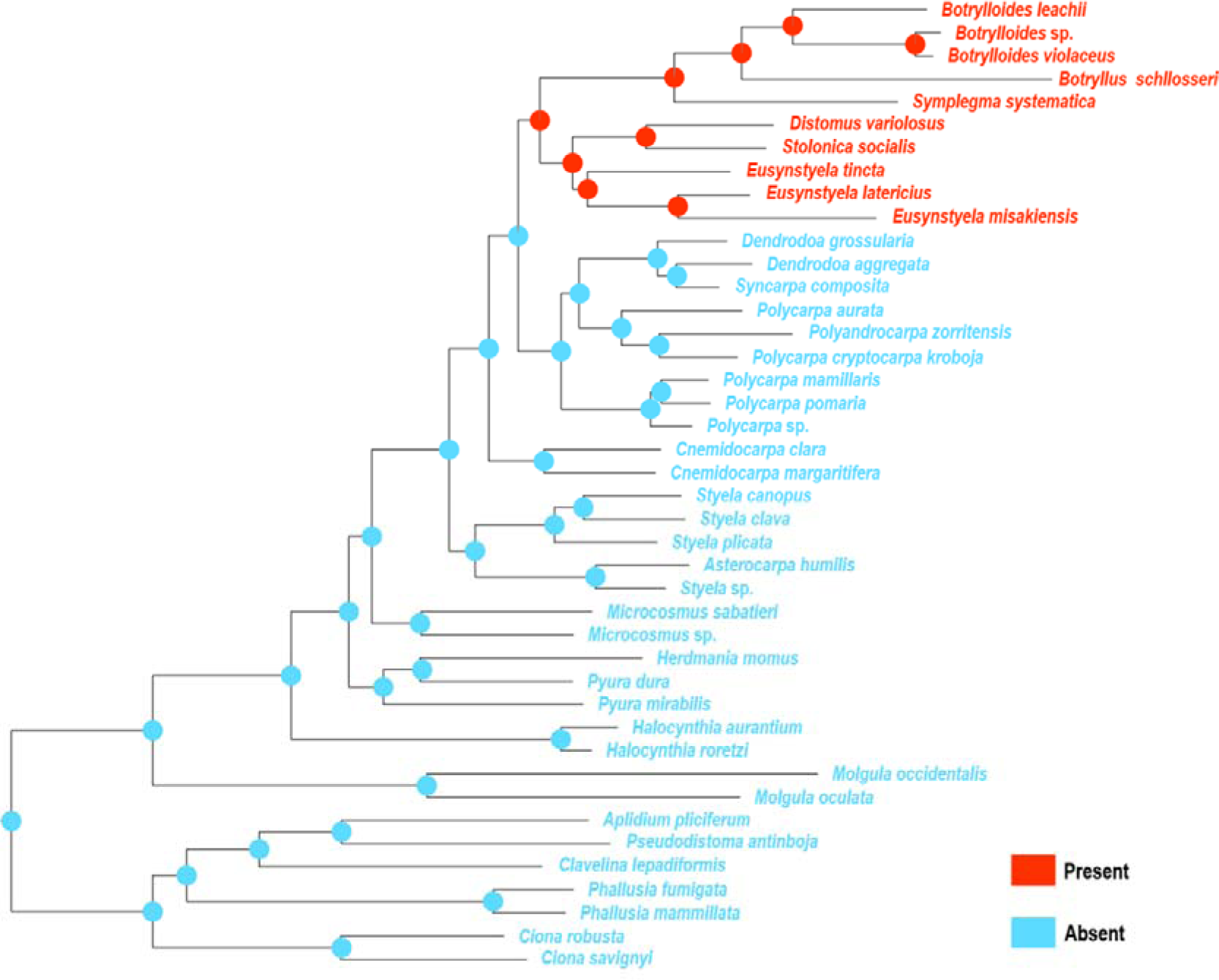
ML tree showing the presence of peribranchial budding. The pie charts indicate EAPs of the characteristics in the nodes.

**Figure 4.**
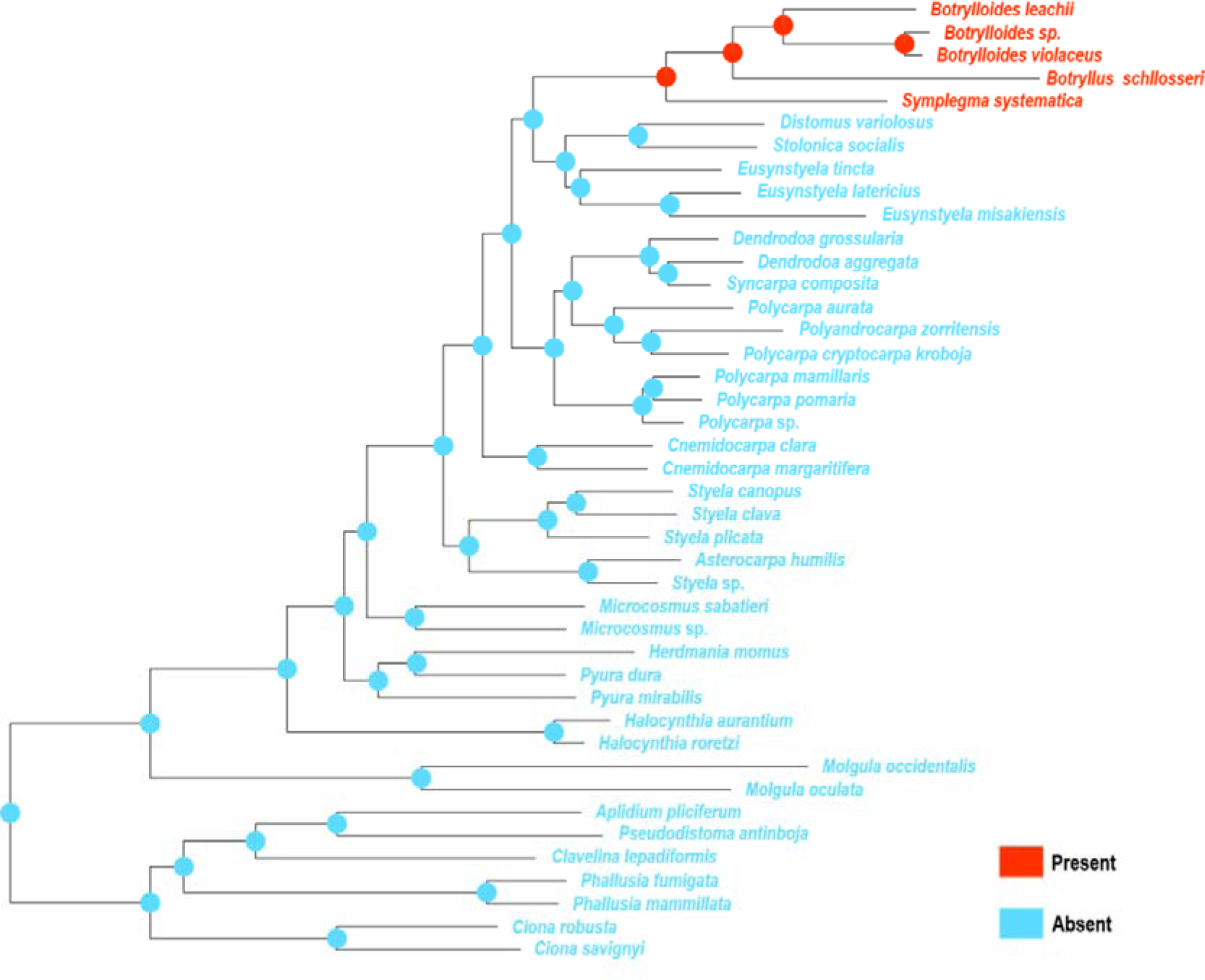
ML tree showing the presence of vascular budding. The pie charts indicate EAPs of the characteristics in the nodes.

**Figure 5.**
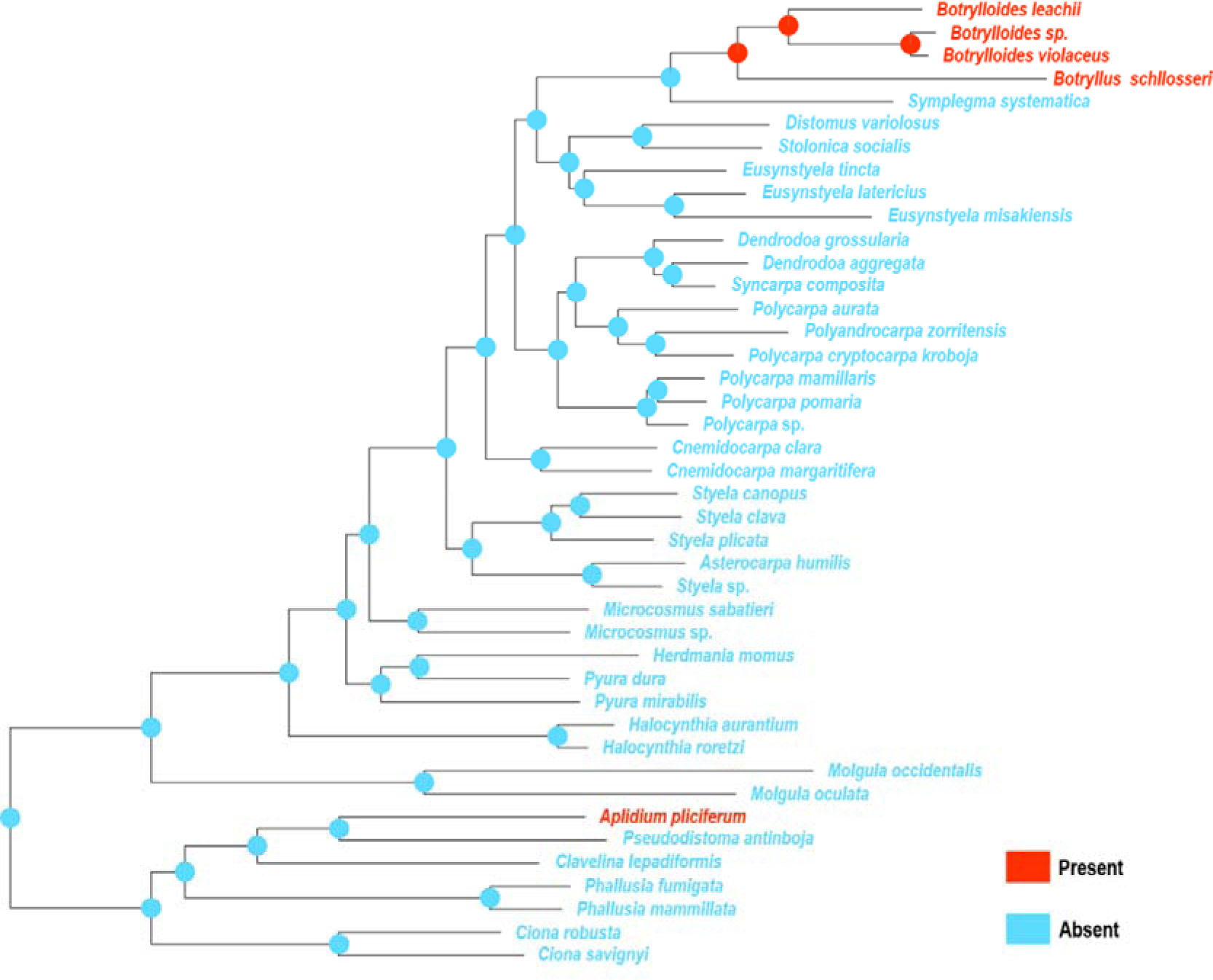
ML tree showing the presence of common cloacal cavity. The pie charts indicate EAPs of the characteristics in the nodes.

**Figure 6.**
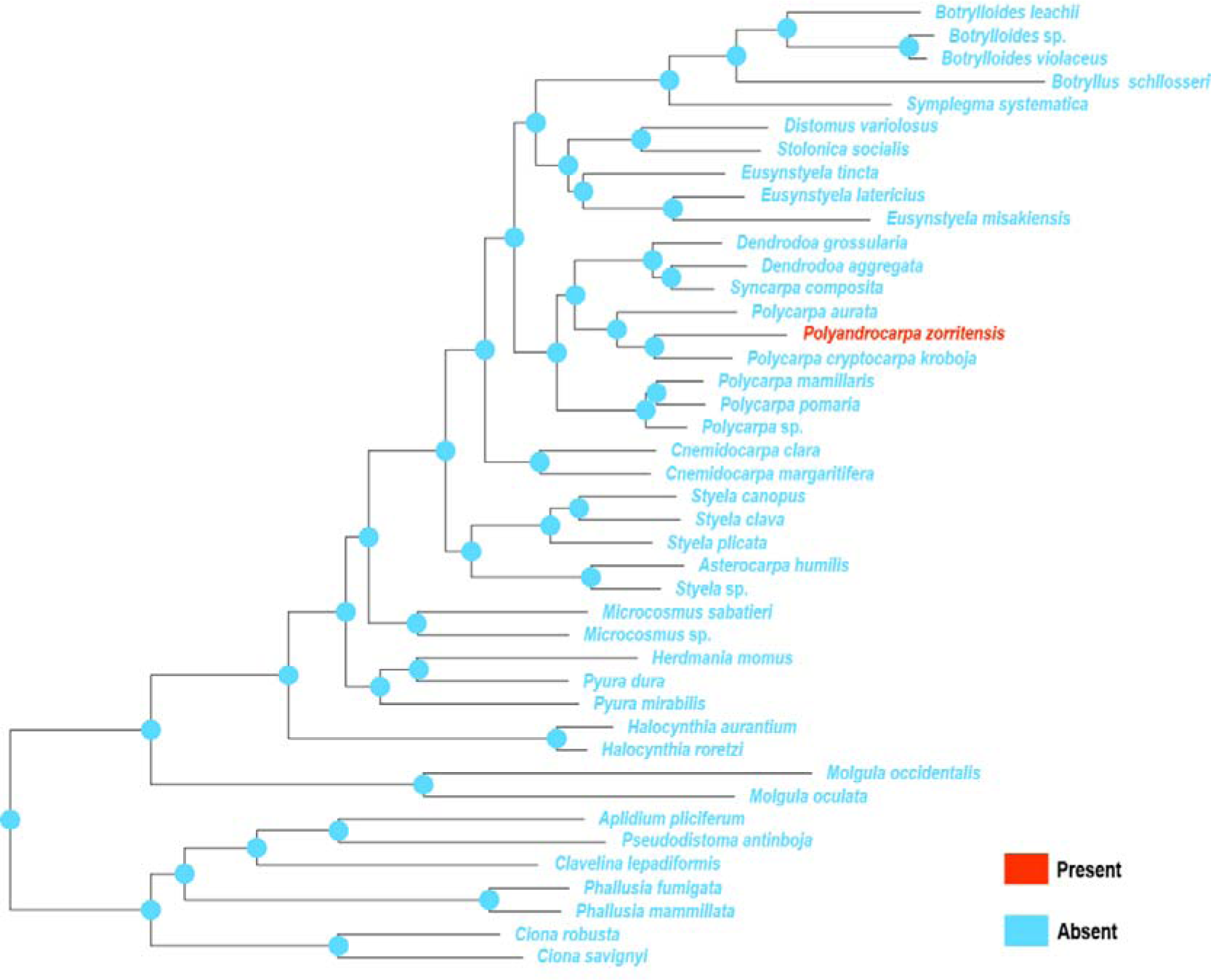
ML tree showing the presence of vasal budding. The pie charts indicate EAPs of the characteristics in the nodes.

**Figure 7.**
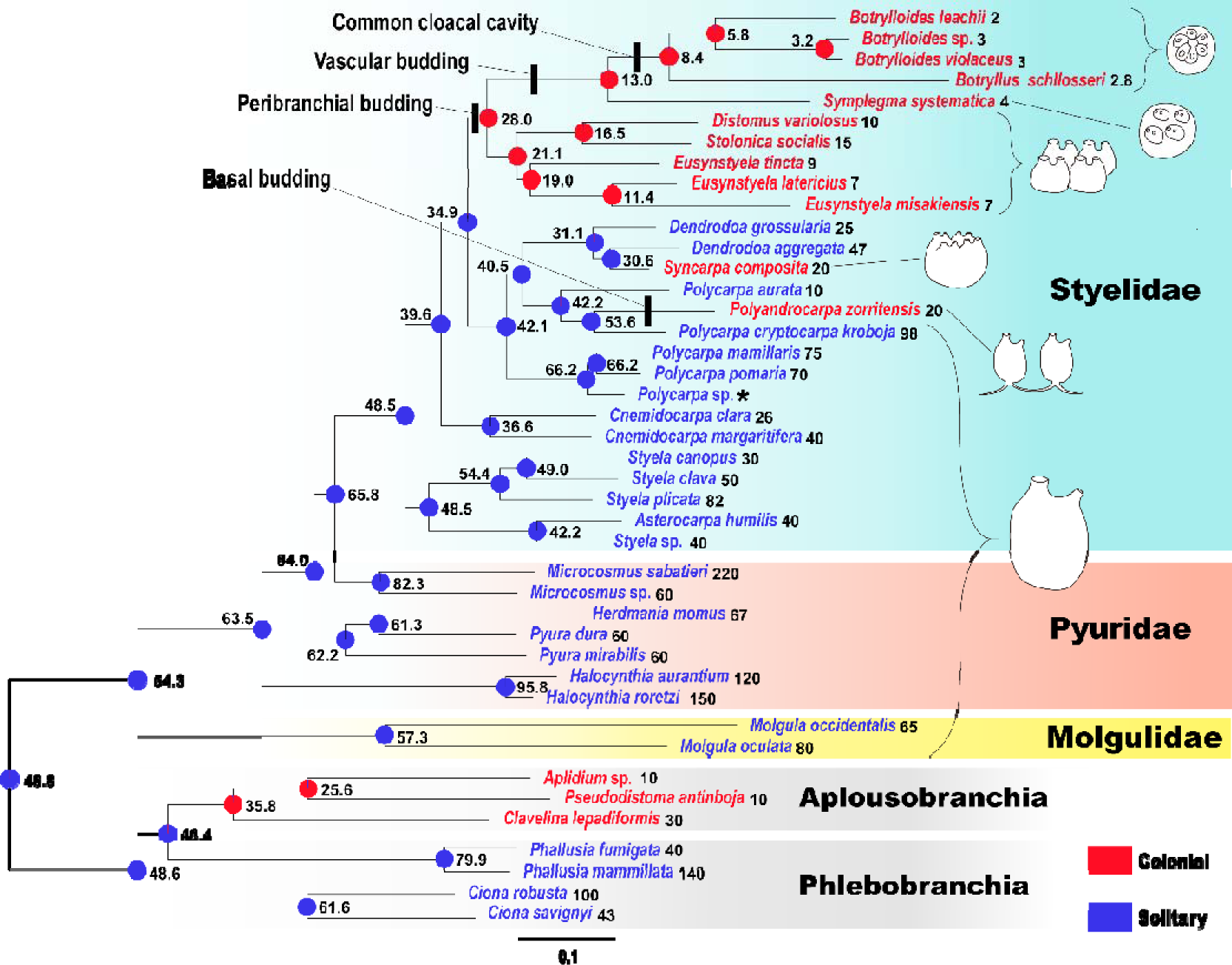
Evolutionary changes of coloniality and body length inferred by ancestral state reconstruction. Pie charts indicate EAPs that an ancestor at each node was colonial/solitary. The numbers show body length of ancestors and OTU; the body length of *Polycarpa* sp. was unknown, so it was indicated by asterisk (*).

### Character correlation

The log marginal likelihoods were −240.18 and −239.98 using complex models in “Test trait correlations: continuous” and “Independent Contrast: Correlation”, respectively, for testing whether there is a correlation between the characteristic states of coloniality and body/zooid length; the log marginal likelihoods with simple models setting the covariance to zero were −242.32 and −242.26, respectively. The log BFs were 4.29 and 4.56, respectively; the values exceeded two, which means positive evidence of correlation between the character states in the two traits.

### Evolutionary direction

The log marginal likelihood based on the entire phylogenetic tree using Model A was −239.80. When using Model B, the value was −240.47. From these two values, the Log BF was calculated to be 1.34. This value is smaller than the threshold value of two, which does not support directionality in trait changes. Based on Clade A, the value using Model A was −109.33; it was −112.35 using Model B. The Log BF was 6.05; as it falls within 5–10, the presence of directionality was strongly supported.

The correlation coefficients between body lengths and branch lengths were 0.01 for solitary forms and −0.85 and colonial forms, respectively. The slopes of the regression lines for these two parameters were ca 2.8 in solitary form (coefficient of determination: R^2^ = 0.0002) and ca. −46.3 in colonial form (R^2^ = 0.7292), respectively (Fig. 8).

**Figure 8.**
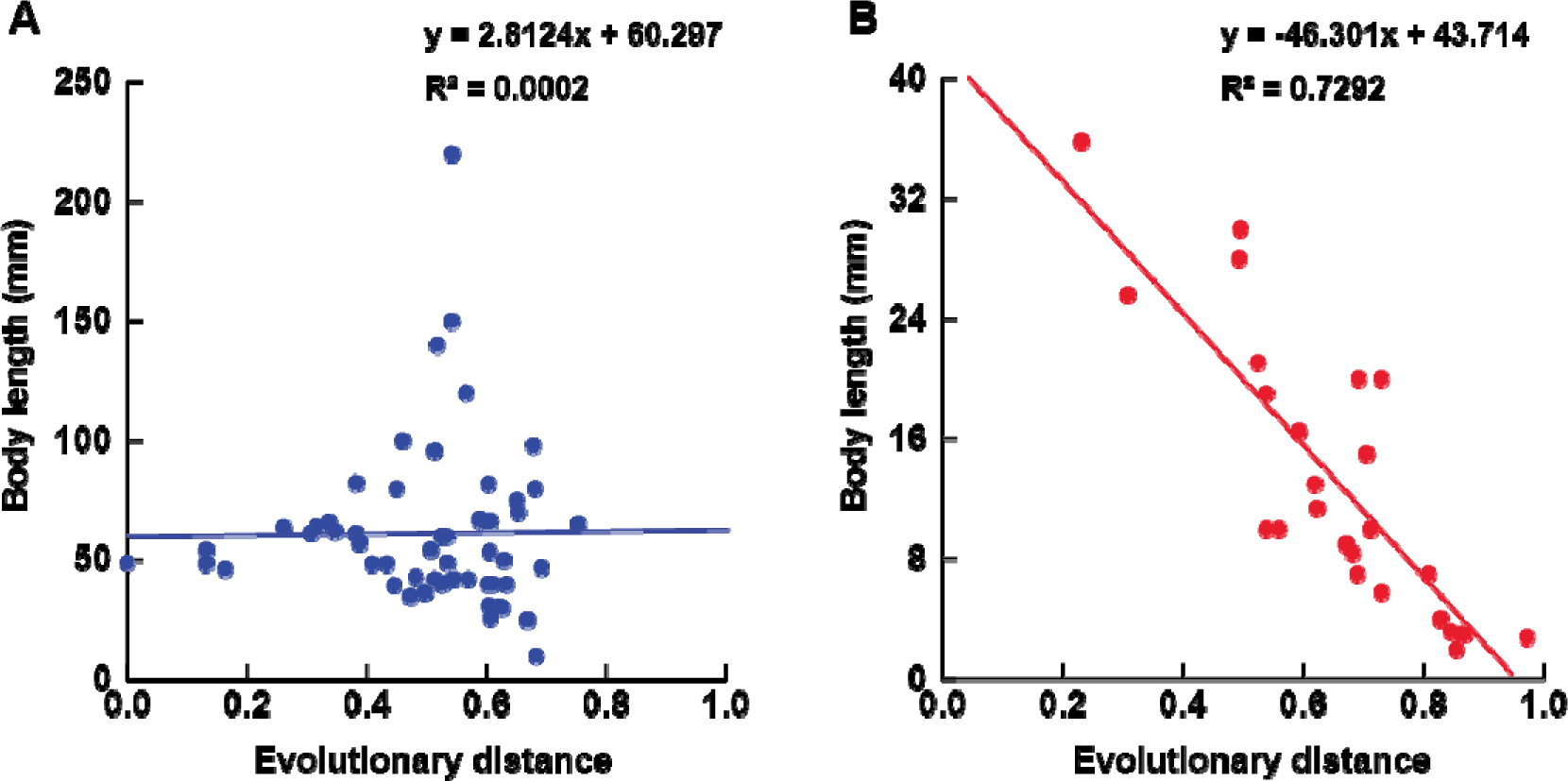
Relationship between body length and evolutionary distance in the OTUs and the nodes of the phylogenomic tree. A, solitary forms (n = 56); B, colonial forms (n = 25)

## Discussion

This study indicates that zooids became gradually smaller once coloniality was acquired (Figs 7, 8). To explain this evolutionary pattern, we hypothesize that zooid miniaturization would shorten the period for daughter zooids to start budding (typically, as an animal’s body size increases, the time it requires to reach maturity also extends; Blueweiss et al. 1978), leading to rapid expansion of the colony over substrates (Fig. 9). The marine substrates represent a significant limiting resource for sessile organisms, with living space being at a premium (Connell 1961; Pequegnat 1964; Dayton 1971; Paine 1971; Jackson 1977, 1985; Hughes 2005; Tyrrell and Byers 2007). Colonial invertebrates achieve this by continuously covering the substrate and exhibiting rapid lateral growth through asexual reproduction with a rate of expansion, which is unattainable through sexual reproduction alone (Jackson 1977).

**Figure 9.**
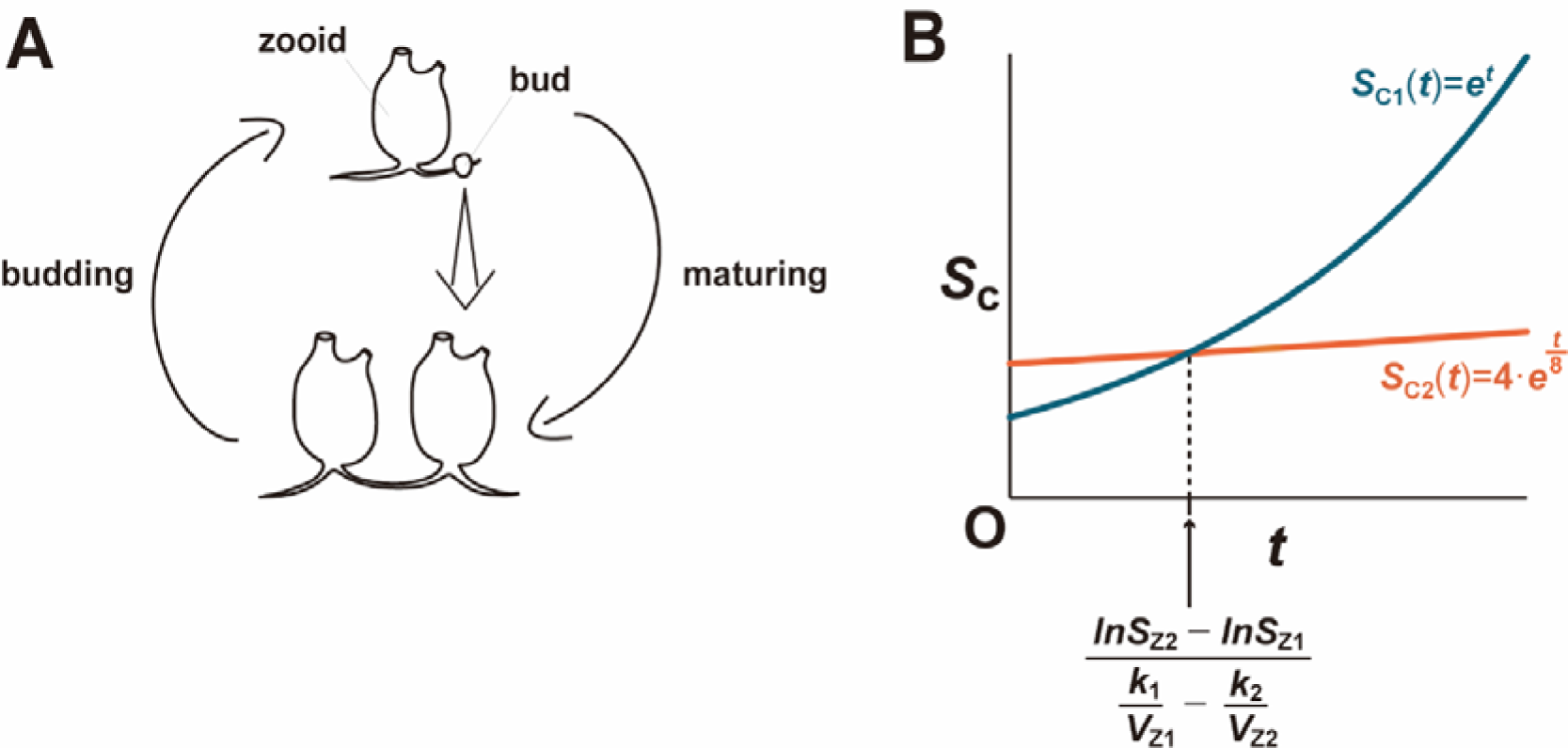
Illustrations of the relationship between zooid size and growth rate of a colony. A, asexual cycle of a budding colony; B, a graph representing two different colonies having similar geometrical structure (*S*_Z_ = 2 or 4; *V*_Z_ = 1 or 8; *k* = 1).

The relationship between the substrate encrusting rate of a colony and the zooid size at maturity can be mathematically described as follows. Let *A*_C_(*t*) be the colony area at time *t*, *V*_Z_ be the zooid volume at maturity, and *k* be the zooid’s volume–growth rate per unit time. The growth rate of colony area per unit time can be expressed by the following equation:

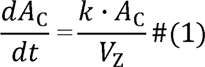

because the area of a colony increases proportionally to the number of zooids in the colony, which is proportional to the colony area. Solving Eq. (1) yields:

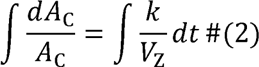

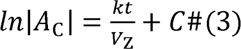

where *C* is an integration constant. Since *A*_C_ > 0, Eq. (3) can be rearranged as:

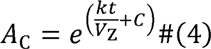

where *e* indicates Euler’s number. Setting *B* = *e^C^* (*B* is a constant), we yield

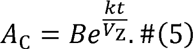

Plugging *t* = 0 in Eq. (5), we obtain:

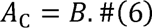

The initial value of the colony area *A*_C_(0) is equal to the area of a single zooid that arises when the larva attaches to the substrate and then metamorphoses. If we denote the zooid area as *A*_Z_, the initial condition *A*_C_(0) equals to *A*_Z_. Plugging this into Eq. (6), we obtain:

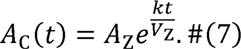

By considering Eq. (7), the competition between two colonies with different zooid volumes and areas can be examined. Let us assume that Colony 1 possesses a larger zooid volume and area compared to Colony 2 (with *A*_Z1_ > *A*_Z2_ > 0 and *V*_Z1_ > *V*_Z2_ > 0), while their respective zooid’s volume growth rates, denoted as *k*_1_ and *k*_2_ (each being a constant). Commencing at *t* = 0, both Colony 1 and Colony 2 originate from a single zooid, where the initial areas can be expressed as *A*_C1_(0) = *A*_Z1_ and *A*_C2_(0) = *A*_Z2_. The value of *t* at which the areas of both colonies become equivalent is determined by the following equation:

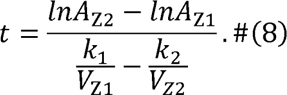

In scenarios where *k*_2_/*V*_Z2_ exceeds *k*_1_/*V*_Z1_, the implication is that Colony 2, with its smaller zooids, will eventually surpass the area of Colony 1, which comprises larger zooids (Fig. 25). Conversely, if *k*_2_/*V*_Z2_ is less than *k*_1_/*V*_Z1_, the resultant negative value of *t* suggests that the area of *A*_C2_ will not overtake that of *A*_C1_. Therefore, this model implies that Colony 1 must accelerate the growth rate of its zooids to achieve a faster colony expansion than Colony 2.

It is important to note here that the model merely discusses the advantages of smaller zooids in colony expansion rates and does not necessarily indicate that colonies with smaller zooids will invariably produce more offspring in subsequent generations. At least three disadvantages with smaller zooids can be conceived: *i*) the zooids would become physically weaker, *ii*) they would be more susceptible to external environmental impacts on the body surface, and *iii*) the number of eggs or sperm per zooid would be reduced. The size of the zooids would be determined by a trade-off with these disadvantages. Moreover, even when the two curves intersect as depicted in Eq. (8) and Fig. 9, the lineage of Colony 1 does not necessarily go extinct if the colony succeeds to form a sufficiently large area to reproduce for the next generation before the time of intersection.

Coloniality was independently acquired at least three times in the evolutionary history of Stolidobranchia (Fig. 7). These results contrast with previous phylogenetic and phylogenomic analyses; acquisition of coloniality was inferred to occur only once (Zeng et al. 2006; Tsagkogeorga et al. 2009; Hasegawa and Kajihara 2019) or at least seven times independently (Pérez-Portela et al. 2009) based on 18S and COI, and at least twice based on transcriptome data (Alié et al. 2018). In addition, the phylogenomic tree, with high support values, supported the view that *Symplegma* would be an intermediate group between other polyzoines and botryllines, an idea earlier proposed based on morphological and immunological evidence (Shirae et al. 1999; Gutierrez and Brown 2017). These results indicate that part of descendants from the last common ancestor of the ascidians that undergo peribranchial budding sequentially acquired vascular budding and common cloacal cavity. The result that the genus *Syncarpa* did not form a clade with other colonial ascidians suggests the potential discovery of a new budding mode or convergent evolution of peribranchial/vasal budding in the future; the budding mode has not been recorded in *Syncarpa* (cf. Redikorzev 1913; Tokioka 1951; Sanamyan 2000; Hasegawa and Kajihara 2019).

In the evolution of coloniality within Stolidobranchia, once traits related to coloniality were acquired, it seemed hard to lose them. The phylogenetic tree did not depict evolutionary pathways reverting from colonial forms back to solitary (Fig. 7); in addition, each budding mode and common cloacal cavity have never been lost after they were acquired. This tendency can also be observed broadly within Tunicata, with some species within the diazonid genus *Rhopalaea* being hypothesized to have undergone an evolutionary loss of budding (Alié et al. 2021). Likewise, very few species in Bryozoa have secondarily acquired solitary forms (Schwaha et al. 2020); no example of reversion from coloniality to solitary in hemichordates (cf. Cannon et al. 2013) and kamptozoans (cf. Fuchs et al. 2010). In contrast, reversal evolution from colonial to solitary forms (and vice versa) is not uncommon in cnidarians (e.g., McFadden et al. 2021).

## Acknowledgements

We would like to express our sincere gratitude for the significant contribution to our sample collections in each field, made by Shoichi Hamano and Hidenori Katsuragawa (Akkeshi Marine Station, Hokkaido University) at Akkeshi Bay; Hisanori Kohtsuka (Misaki Marine Biological Station, the University of Tokyo) at Misaki; Masashi Fukuoka, Naoto Jimi, Misato Sako, and Maki Shirae-Kurabayashi (Sugashima Marine Biological Laboratory, Nagoya University) around Sugashima Island; Takeo Sugimoto (South Wakayama Fishery Cooperative), Natsumi Hookabe (Japan Agency for Marine-Earth Science and Technology, JAMSTEC), Cati (Association for the Protection of the Sea and Marine Life, *Umimorinchu*), Ayumi Mizutani, Takashi Okada, and Naofumi Ueda (Dive KOOZA) in Kushimoto and Koza; Hiroshi Kuriyama (Miyako Island Fishery Cooperative), Hiroki Iranami (Irabu Island Fishery Cooperative), Hiroshi Yonamine (Ikema Island Fishery Cooperative), Yosuke Kamata, Manami Kimura, Junko Watanabe (Aquastar Miyakojima), Yuki Kita, Hiroki Matsushita, Shoki Shiraki, and Aoi Tsuyuki (Hokkaido University) around Irabu Island and Miyako Island; Bar Gabso, Lion Novak (Tel-Aviv University), Nahum Sela (Interuniversity Institute for Marine Sciences), and a volunteer diver at Interuniversity Institute for Marine Sciences, Shlomi Levi, in Eilat, Israel. Without the invaluable assistance given to NH provided by Gil Koplovitz (Interuniversity Institute for Marine Sciences) and Gal Vered (Tel-Aviv University) during NH’s lab works at Interuniversity Institute for Marine Sciences, the goals of this research would not have been realized. A precious photograph of *Polyandrocarpa zorittensis* given by Teruaki Nishikawa improve the figure in this study. Our sincere thanks are offered for the academic support provided by Analytical Center of Suntory Foundation for Life Sciences Bioorganic Research Institute, SUNBOR in conducting RNA-seq analysis. We would like to extend our heartful thanks to Kei Kitahata, whose technical insights were a cornerstone in the creation of the scripts that enabled smooth progress in our phylogenomic analyses. We are deeply grateful to all the individuals who generously contributed to NH through the academic crowdfunding platform “academist”; special thanks go to Shunji Furukuma, Naoki Hayashi, Miyuki Honda, Hitoki Horie, Sho Hosotani, Yoshiki Iwai, Nami Kenmotsu, Moe, Takehiro Nakamura, Ryoma Nishikawa, Yuichi Sasaki, Tatsuya Shimoyama, Makoto Taniguchi, Daiki Wakita, Takaaki Yonekura, and many others. NH was granted financial support from JST SPRING, with Grant Number JPMASP2119.

